# Molecular prosthetics and CFTR modulators additively increase secretory HCO_3_^−^ flux in cystic fibrosis airway epithelia

**DOI:** 10.1101/2025.06.18.660463

**Authors:** Nohemy Celis, Danforth P. Miller, Thomas E. Tarara, Jeffry G. Weers, Ian M. Thornell, Michael J. Welsh, Martin D. Burke

## Abstract

Cystic Fibrosis (CF) is caused by loss-of-function mutations in the gene encoding the cystic fibrosis transmembrane conductance regulator (CFTR), an anion channel predominantly expressed on the apical membrane of epithelial cells. Reduced Cl^−^ and HCO_3_^−^ secretion due to dysfunctional CFTR results in a decrease in lung function and is the leading cause of morbidity in individuals with CF. Recent therapies, known as highly effective CFTR modulator therapy (HEMT), help improve the lung function in individuals with specific CF-causing mutations by enhancing the folding, trafficking, and gating of CFTR. However, variability in HEMT responsiveness leads to sub-optimal clinical outcomes in some people with CF undergoing modulator therapy. A potential strategy is to complement their function with a CFTR-independent mechanism. One possibility is the use of ion channel-forming small molecules such as amphotericin B, which has shown promise in restoring function and host defenses in CF airway disease models. Amphotericin B functions as a molecular prosthetic for CFTR and may thus complement CFTR modulators. Here we show that co-treatment of CF airway epithelia with HEMT and amphotericin B results in greater increases in both HCO_3_^−^ secretory flux and ASL pH compared to treatment with either agent alone. These findings suggest that co-administration of CFTR modulators and molecular prosthetics may provide additive therapeutic benefits for individuals with CF.

## INTRODUCTION

Cystic Fibrosis (CF) is a genetic disease characterized by severe respiratory complications caused by mutations in the *CFTR* gene, which encodes the cystic fibrosis transmembrane conductance regulator (CFTR) protein^1^. The CFTR protein is an anion channel primarily located on the apical membrane of epithelial cells and is critical for maintaining hydration, ion balance, and pH of the airway surface liquid (ASL)^2^. Basolateral pumps such as the Na^+^ /K^+^ -ATPase establish an electrochemical gradient that facilitates HCO_3_^−^ entry into epithelial cells, providing a driving force for transepithelial HCO_3_^−^ secretion through CFTR (Figure 1A)^3^. Loss-of-function mutations in CFTR disrupt this process by reducing HCO_3_^−^secretion and lowering ASL pH. The resulting acidic environment impairs host defenses, promotes the accumulation of thick, viscous mucus, and compromises mucociliary clearance leading to persistent infection and inflammation in the lungs (Figure 1B)^4,5^.

**Figure 1.**
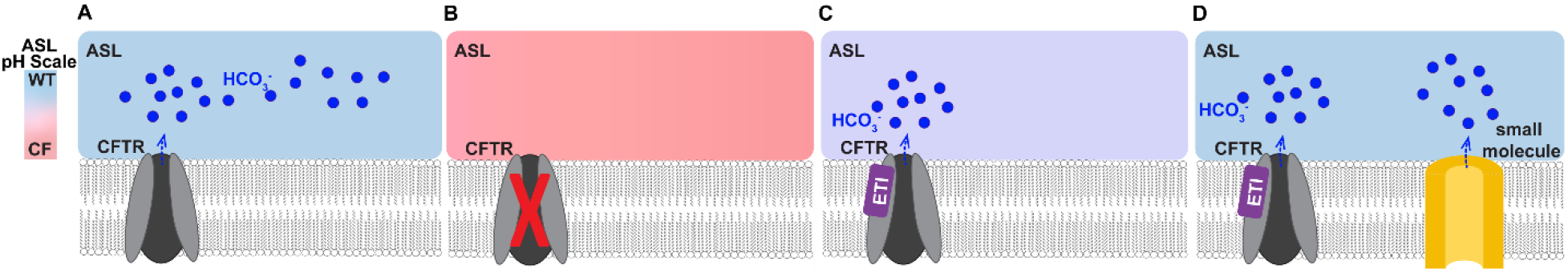
Small molecule ion channels complement the function of CFTR modulators in cystic fibrosis airway epithelia. **A**) The apical anion channel CFTR regulates HCO_3_^−^ efflux in epithelial cells. **B**) Mutations in CFTR result in complete or partial loss of HCO_3_^−^ secretion, impairing the ASL pH, viscosity and antibacterial activity. **C**) CFTR modulators (Elexacaftor (E), Tezacaftor (T), and Ivacaftor (I)) partially restore CFTR function. **D**) We hypothesize that co-treatment of CFTR modulator treated airway epithelia with small molecule-based Amphotericin B ion channels will result in more secretory HCO_3_^−^ flux.

The development of CFTR modulators, such as Elexacaftor/Tezacaftor/Ivacaftor (ETI), have led to remarkable advances in CF therapy by improving CFTR function in individuals with responsive mutations (Figure 1C)^6–8^. However, the improvement in transepithelial Cl^−^ and HCO_3_^−^secretion and the clinical response to ETI varies across individual patients, even those with the same mutations^9–12^. Given that HCO_3_^−^secretion plays a key role in airway host defenses^13–17^, we reasoned that an alternative protein-agnostic pathway could increase HCO_3_^−^secretion by complementing the function of current CFTR modulators. One promising approach is the use of amphotericin B (AmB), a small molecule that can form ion channels and functionally complement CFTR as a molecular prosthetic (Figure 1D) ^18^. Unlike CFTR modulators, molecular prosthetics function independent of CFTR. Pre-clinical studies have demonstrated the potential of molecular prosthetics to restore ion transport, increase ASL pH, improve ASL hydration, decrease ASL viscosity, and increase ASL antibacterial activity in cultured CF airway epithelia^18^. Additionally, early clinical studies showed that, compared to vehicle control, AmB nasal perfusion in a zero Cl^−^solution caused a more negative nasal potential difference in people with CF not on CFTR modulators^19^. These potential differences were similar to those observed in early clinical studies with Ivacaftor^19–21^. With these promising results, AmB was formulated as an inhalable dry powder drug-device combination product (CM001) and is currently being evaluated in Phase 1 clinical trials^22–24^. The lead formulation of CM001 (ABCI-003), comprising lipid-coated crystals of the molecular prosthetic, was utilized for the cell-based assays described herein.

In this study, we tested whether co-treatment of CF airway epithelia with ETI and AmB can lead to an additive increase in HCO_3_^−^ secretory flux. Our findings reveal that co- treatment of CF airway epithelia with ETI and AmB additively increases HCO_3_^−^ secretory flux to the ASL and thereby increases ASL pH.

## RESULTS

A concentration-response analysis with ETI on cultured CF airway epithelia was performed to assess HCO_3_^−^secretory flux and ASL pH. As expected, ETI (E=1µM, T=1µM, I=0.33µM) increased transepithelial H^14^CO_3_^−^secretion and ASL pH (Fig. 2A-C). Notably, there were no further increases in HCO_3_^−^ secretory flux or ASL pH upon addition of up to 10-fold higher concentrations of CFTR modulators relative to the lowest concentration of ETI used for these experiments (Figure 2A-C).

**Figure 2.**
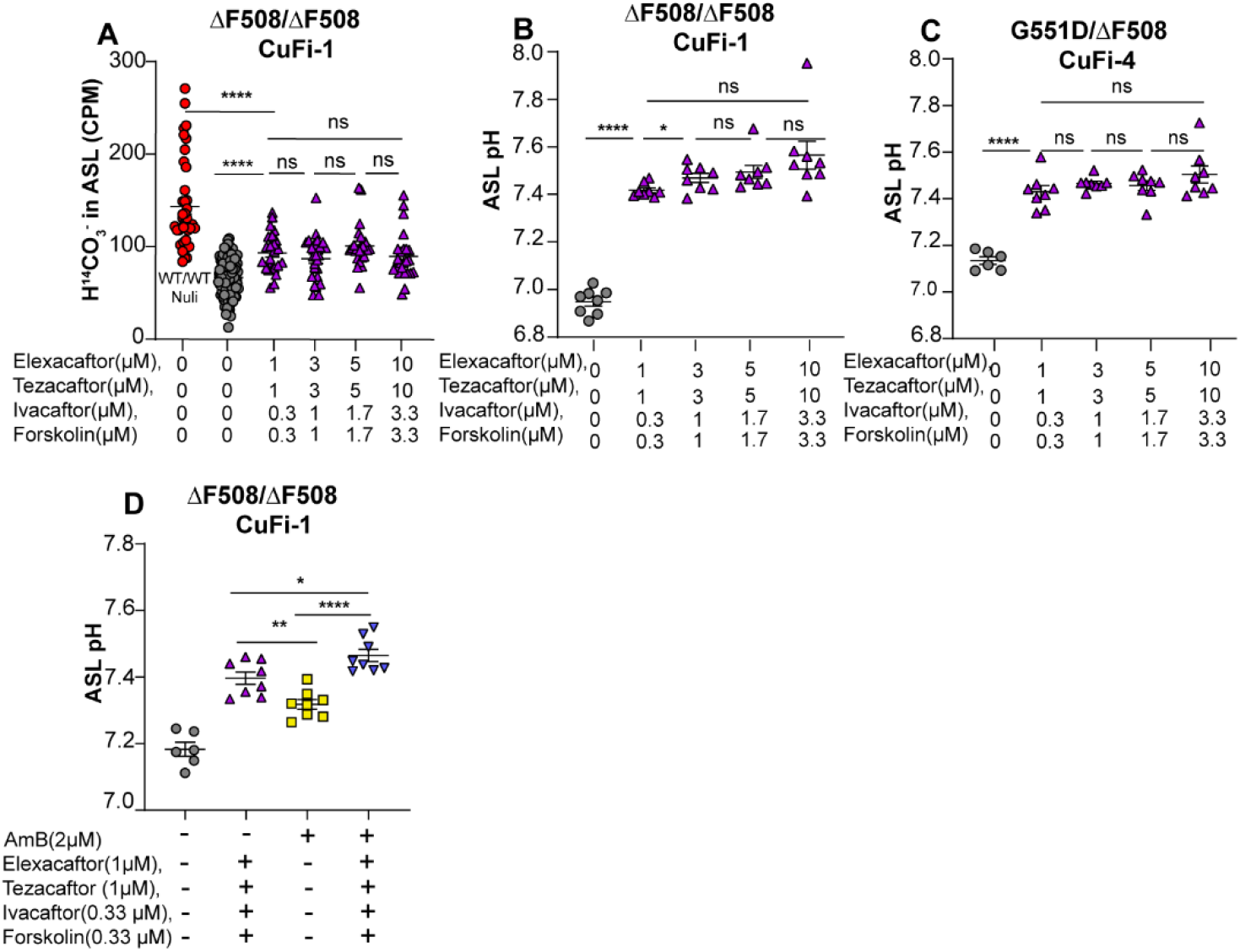
The ceiling effect of CFTR modulators. **A-C**) Incubation with increasing concentrations of CFTR- modulators (ETI) (measurements taken 48 hrs. after modulator treatment) does not lead to further increases in secretory H^14^CO ^−^ flux to the ASL or increases in ASL pH in CF airway epithelia in comparison to the lowest dose of ETI tested here. **D**) 48 hour treatment with either ETI/forskolin or AmB increases ASL pH in CuFi-1 epithelia ; Additive effects are observed when these two treatments are combined. Student T-test used for statistical analysis: ns not significant, *P≤0.05, **P ≤0.01, ***P ≤0.001, **** P ≤0.0001.

To evaluate whether AmB based channels led to further increases in ASL pH in ETI treated airway epithelia, we co-incubated cultured CF airway epithelia with ETI, AmB, or both (Figure 2D). As expected, the addition of CFTR modulators to CuFi-1 (ΔF508/ΔF508) epithelia led to an increase in ASL pH. CuFi-1 epithelia treated with AmB also led to increases in ASL pH, albeit to a somewhat lower extent than CFTR modulators. Notably, treatment with both CFTR modulators and AmB led to a significant increase in ASL pH relative to either treatment alone. Further studies with an Ussing chamber confirmed that AmB-based channels could still form in ETI treated CF airway epithelia (Figure S1A-C). This led to the conclusion that the complementary mechanisms of both classes of small molecules resulted in the observed additive increases in ASL pH.

Given that AmB-based channels retain activity in ETI-treated CF airway epithelia and contribute to additive increases in ASL pH, we conducted an extended set of combination studies using ABCI-003. We first tested whether AmB ion channels in the form of ABCI-003 increased HCO_3_^−^secretory flux in modulator-treated cultured CF airway epithelia. Co-treatment of CuFi-4 (G551D/ΔF508) epithelia with both modulators and ABCI-003 for 48-hours led to a substantial increase in H^14^CO^3-^secretory flux (Figure 3A- C). In addition, we found that the inhibition of Na^+^ /K^+^ -ATPase with ouabain stopped the secretion of H^14^CO_3_^−^. These findings suggest that, similar to CFTR, HCO_3_ ^−^ secretion through small molecule ion channels is influenced by the electrochemical gradient maintained by the Na^+^ /K^+^ -ATPase.

**Figure 3.**
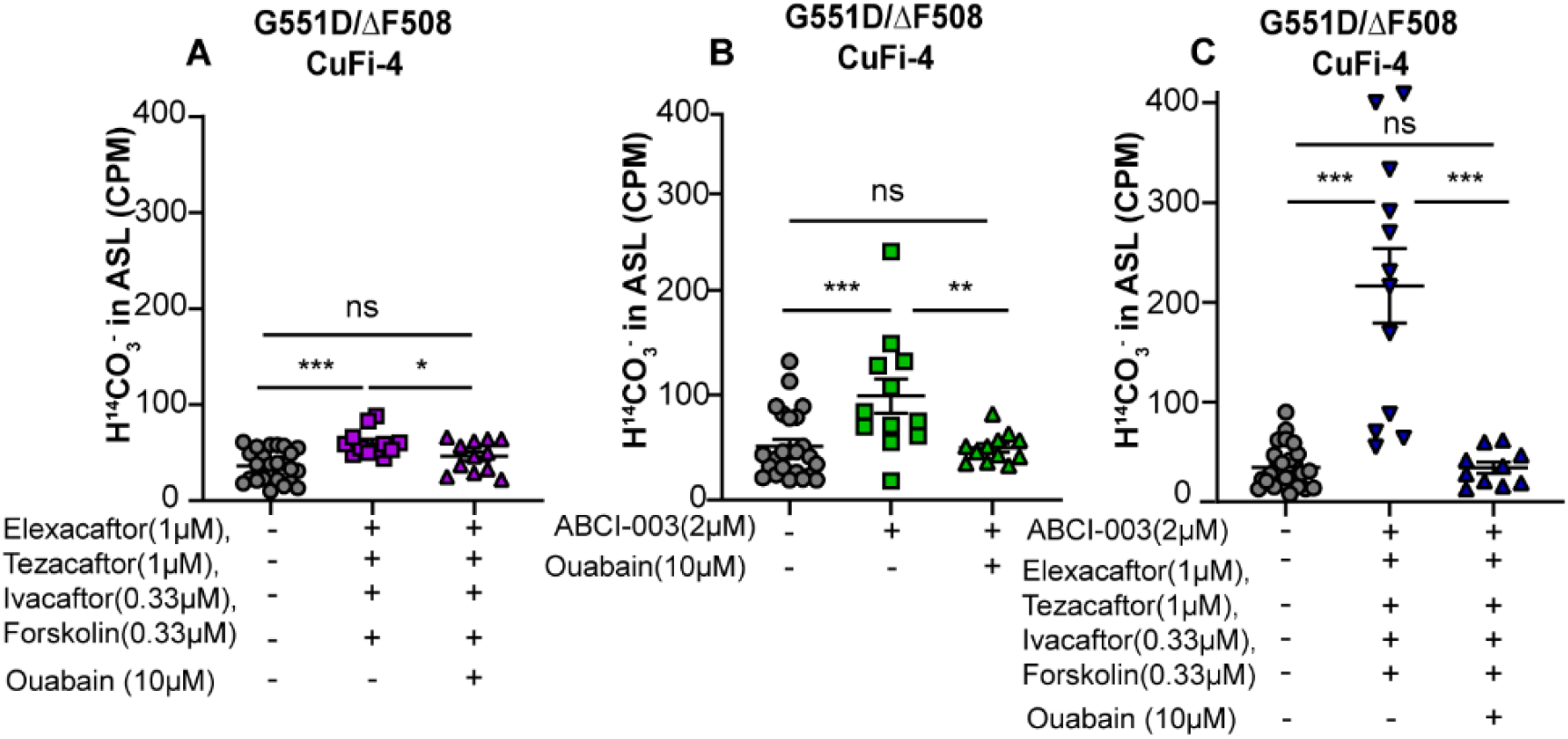
HCO_3_^−^ secretory flux in CF airway epithelia is powered by the basolateral Na+/K+ ATPase. **A-B**) H^14^CO_3_^−^ secretory flux in CuFi-4 cultured lung epithelial cells post-48 hrs. by treatment with CFTR-modulators or ABCI-003. **C**) Co-treatment of CFTR modulator treated airway epithelia with ABCI-003 leads to an additive H^14^CO_3_^−^ efflux (**A-C**) Inhibition of the Na+/K+ ATPase with ouabain ceases apical secretory_3_ H^14^CO_3_^−^ flux through small molecule ion channels (ABCI-003) and rescued CFTR. Student T-test used for statistical analysis: ns not significant, *P≤0.05, **P ≤0.01, ***P ≤0.001, **** P ≤0.0001.

We then exposed cultured CF airway epithelia for 48 hours to different concentrations of CFTR modulators while keeping the ABCI-003 concentration the same to assess ASL pH and secretory H^14^CO_3_^−^flux (Fig. 4A-H). Notably, ABCI-003 increases ASL pH and promotes H^14^CO_3_^−^ secretory flux to a similar extent as CFTR modulators (Figure 4A-H). In addition, additive effects with ABCI-003 addition were also observed when CF airway epithelia were treated with a low and higher concentration of CFTR modulators, highlighting complementarity between these mechanisms.

**Figure 4.**
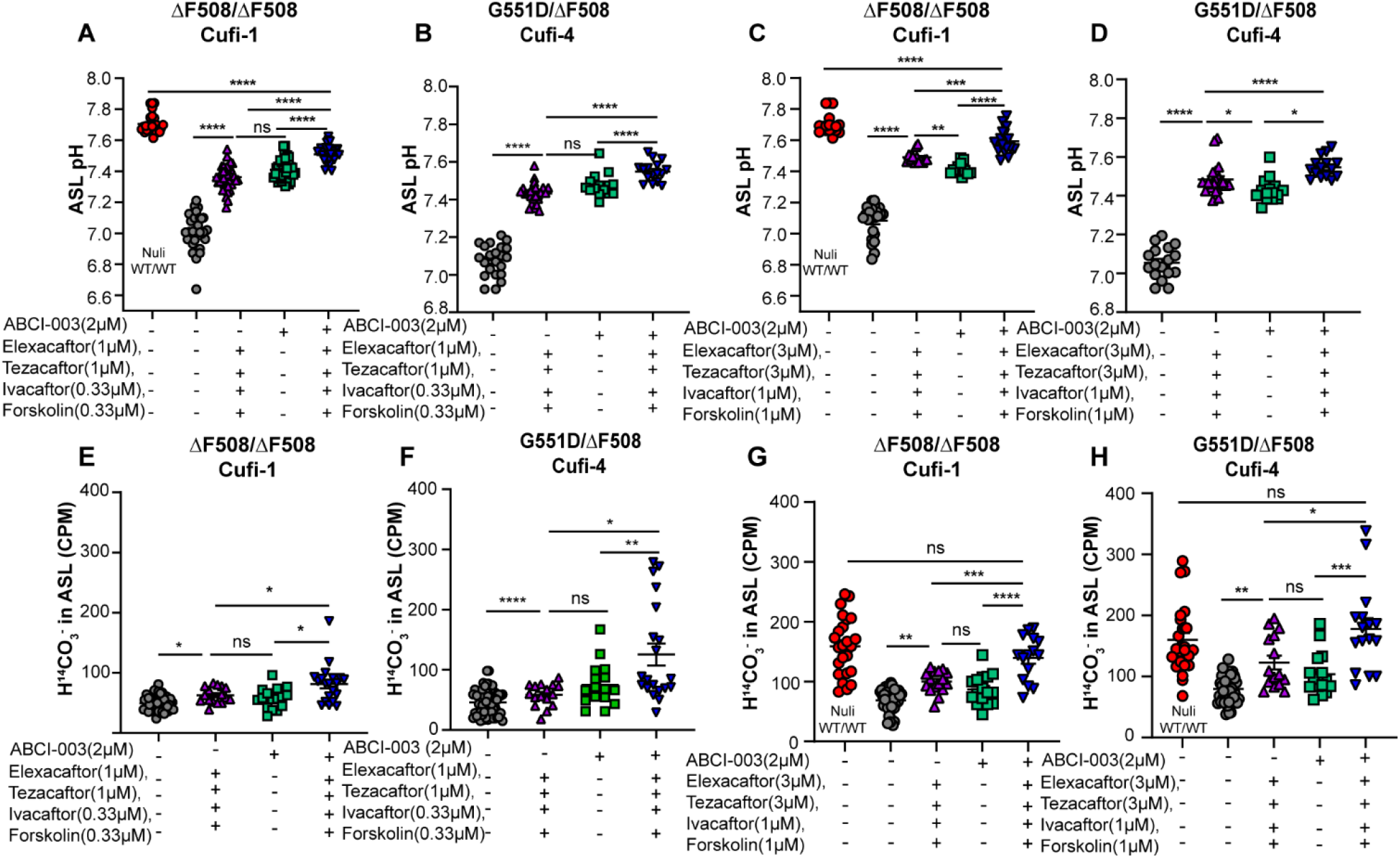
Molecular prosthetics and CFTR-modulators work in complement to additively increase ASL pH and H^14^CO_3_^−^ efflux in CF airway epithelial cells. (**A-B**) Effect of ETI and ABCI-003 on ASL pH post-48 hrs. incubation in CuFi-1 and CuFi-4 epithelia. (**C-D**) Effect of higher ETI concentrations and a fixed ABCI-003 concentration of 2µM on ASL pH in CuFi-1 and CuFi-4 epithelia. (**E-F**) CuFi-1 and CuFi-4 epithelia H^14^CO_3_^−^ secretory flux from basolateral to the ASL post-48hrs. incubation with ETI, and ABCI-003 (**G-H**) CuFi-1 and CuFi-4 epithelia H^14^CO_3_^−^ secretory flux from basolateral to the ASL post-48hrs. incubation with increased ETI concentrations and a fixed ABCI-003 concentration at 2µM. ns not significant, *P≤0.05, **P ≤0.01, ***P ≤0.001, ****P ≤0.0001.

## DISCUSSION

The effect of pH changes due to altered HCO_3_^−^ secretion in the ASL has repeatedly been demonstrated to be important for antibacterial activity, mucociliary clearance, and mucin expansion _13,14,14–18,25–28_. We compare for the first time the efficacy of the clinical formulation ABCI-003 to highly effective CFTR modulator therapy. We show that ABCI- 003 restores ASL pH and HCO_3_^−^ efflux in CF preclinical models to a similar capacity as HEMT. These results suggest that ABCI-003 on its own could potentially benefit people with CF who either must discontinue HEMT due to poor tolerability^8,29,30^ or can potentially benefit those who do not have CFTR modulator responsive mutations. Additionally, our data suggest that the mechanisms of CFTR modulators and molecular prosthetics complement one another to increase secretory HCO_3_ ^−^ flux to the ASL of CF airway epithelia. This strategy may potentially benefit some individuals currently on HEMT experiencing adverse events.^8,29,30^ In such cases, co-administration of ABCI-003 might be beneficial by enabling a reduced ETI dose without sacrificing efficacy. Collectively, these results suggest the potential for additive benefit of molecular prosthetics and CFTR modulators in improving airway host defenses in people with CF.

## METHODS

*Small-diameter NuLi and CuFi cultured epithelia were used for these experiments (0.33 cm^2^)*

### Compound preparation and incubation methods

#### Constituents of the dry powder formulation of Amphotericin B

The drug product, CM001 (amphotericin B inhalation powder), is a drug-device combination comprising a dry powder formulation of Amphotericin B that is administered with a portable, unit-dose, capsule-based dry powder inhaler (DPI). The drug constituent part comprises wet-milled crystals of amphotericin B coated with a porous shell of phospholipids, cholesterol, and calcium chloride. The lead formulation (ABCI-003) is spray-dried from an emulsion-based feedstock and contains 15.8% w/w Amphotericin B. The device constituent part is the ultrahigh resistance variant of the RS01 DPI (Plastiape S.p.A., Osnago, Italy). The lead formulation of CMABCI-003 was utilized for the cell-based assays described^23,24,31–33^.

#### ABCI-003 sample preparation

A target amount of ABCI-003 was added into a 1.5 mL HPLC vial and suspended in 100 μL of FC-72 (PFC). The absorbance of the stock was measured using the cuvette function of a NanoDrop OneC at 406 nm. The extinction coefficient of AmB (164000 M^-1^*cm^-1^) was further used to calculate the concentration of the stock solution. Then, the desired concentrations of ABCI-003 were prepared at 2 μM in a final sample volume of 200 μL of PFC. The samples were vortexed and sonicated for 1 minute prior to the apical addition of 20 μL of sample per cultured lung epithelia well. PFC is used since it readily evaporates and leaves behind the ABCI-003 powder.

#### Modulator(s) sample preparation

Elexacaftor (VX-445), Tezacaftor (VX-661), and Ivacaftor (VX-770) were purchased from Selleckchem as a 5 mg powder form and solubilized in DMSO to prepare 10 mM stock concentrations. Forskolin was purchased from Sigma-Aldrich as a 10 mg solid and solubilized to prepare a 10 mM stock solution in DMSO. 1 mM stock solutions were prepared for each compound listed by obtaining 100 μL from the 10 mM stocks and added into 900 μL of DMSO. 550 μL of fresh 1:1 DMEM:Ham’s F-12, supplemented with 2% v/v Ultroser G (Crescent Chemical) plate media was added to 24 well plates followed by Elexacaftor/Ivacaftor/Tezacaftor and forskolin addition at the desired volume for the experimental modulator concentrations (Elexacaftor/Tezacaftor range: 1 μM-10 μM, Ivacaftor range: 0.33 μM-3.67 μM) and the CFTR activator forskolin range: 0.33 μM-3.67 μM. After the addition of ABCI-003, modulator(s), or both, the cells were incubated at 37° in 5% CO_2_ for 48-hours.

### ASL pH determination-48 hours post compound addition

At least one hour prior to imaging, the Zeiss LSM880 confocal microscope was turned on and set to 5% CO_2_ at 37°C. The excitation wavelength was 514 nm and emission wavelengths were set at a range of: 578-608 nm (channel 1) and 633-663 nm (channel 2). One hour prior to imaging, 2.5 μL of SNARF-dextran in Dulbecco’s PBS (DPBS) was added onto the apical side of the cultured lung epithelia. The cell membranes from each experimental group were then imaged using a 40x objective with water immersion for cell line cultures on the objective lens.

### H^14^CO_3_^−^efflux 48-hours post compound addition

^14^C-labelled sodium bicarbonate was obtained as a sterile 35.7 mM aqueous solution pH 9.5 from PerkinElmer. 5 μL of a 2.8 mM H^14^CO_3_^−^ stock solution in USG medium was added to the basolateral medium. The lung epithelial cells were then incubated in 5% CO_2_ at 37 °C for 1 hr. After 1 hr, the apical membrane was immediately washed with 200 μL of PBS, 150 μL of the wash was collected and added into 3 mL of Opti-Fluor (a high flash point LSC-cocktail for aqueous samples, Thermo Scientific). Additionally, 150 μL of PBS and 150 μL of basolateral media were also added into 3 mL of Opti-Fluor and analyzed by scintillation liquid counting.

### Statistical Analysis

All statistical analyses were performed with GraphPad Prism software. Unpaired t-test with Gaussian distribution and multiple comparisons test were used for quantifying statistical differences between sample groups. Statistical designations: NS not significant, *P ≤ 0.05, **P ≤ 0.01, ***P ≤ 0.001, and ****P ≤ 0.0001.

## ASSOCIATED CONTENT

### Supporting Information

The Supporting Information can be found on the bioRxiv website. Supporting data for Ussing chamber studies.

## AUTHOR INFORMATION

### Notes

The University of Illinois has filed patent applications related to the use of amphotericin B as a molecular prosthetic for CFTR, additive effects with CFTR modulators, and the dry powder formulation CM001. These patents have been licensed to cystetic Medicines, a biotech company for which MDB and MJW are scientific founders, consultants, and shareholders.

## ACKNOWLEDGEMENTS

We are grateful for cystetic Medicines (C0673 Studies with Amphotericin B Cystetic for Inhalation) and the NIH (MIRA R35GM118185 to MDB) for funding this work. We thank the NIH funding support for N.C a Predoctoral Fellow (5T32 GM136629-04) who was also partially supported by the UIUC Peixin He and Xiaoming Chen PhD4 Graduate Fellowship.

