## Supplemental Figure S1 A-C for "Molecular prosthetics and CFTR modulators additively increase secretory HCO_3_^−^ flux in cystic fibrosis airway epithelia"

**Table of Contents**

**S2** Fig. S1. Short circuit current changes from an Ussing chamber showing that AmB permeabilizes CFTR modulator treated CuFi-4 (G551D/ΔF508) airway epithelia. Method section for Ussing chamber studies.

### Supplemental Figures

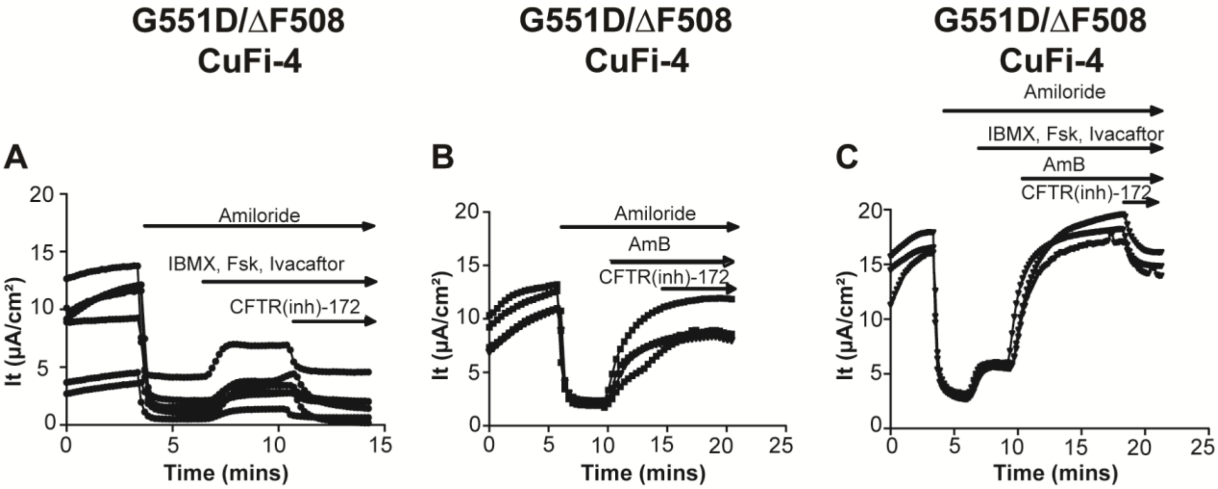


**Fig.S1** AmB permeabilizes CFTR modulator treated CuFi-4 (G551D/ΔF508) airway epithelia. A) Electrocurrent change ($I_t$) of CuFi-4 airway epithelia post-48 hours incubation with Elexacaftor (1μM), Tezacaftor (1μM), and Ivacaftor (0.33μM ) following inhibtion of the sodium channel ENAC with Amiloride (100μM), and activation of CFTR with the phosphodiesterase inhibitor, protein kinase A activator (IBMX-100μM and Forskolin(Fsk)-10μM) and Ivacaftor (10μM). B) $I_t$ increases following addition of small molecule ion channels formed with amphotericin B (AmB-10μM) that are insensitive to inhibtion by the CFTR inhibitor CFTR(inh)-172 (10μM ). C) Activation of CFTR with IBMX (100μM) , Fsk (10 μM), and Ivacaftor (10μM) increases $I_t$ of CuFi-4 airway epithelia post-48 hour incubation with Elexacaftor, Tezacaftor, and Ivacaftor as before. Follow-up addition of AmB (10μM) shows $I_t$ increases, validating small molecule ion channel formation in the presence of modulator rescued CFTR.

**Method**

**Ussing chamber electrophysiology measurements.** Apical and basolateral ringer solutions consisting of: 120mM sodium gluconate /120 mM NaCl , 25 mM NaHCO3, 5 mM KCl, 2 mM CaCl2, 1.2 mM MgCl2, 10mM C6H12O6 and 13.75 mM of NaH2PO4 at pH 7.4 were added to each side of the chamber and maintained at 37 °C while gassed with compressed air. The epithelial Na+ channel (ENaC) was inhibited by a 100 μM addition of Amiloride to the apical side to achieve a baseline for permeabilization/CFTR activity. Forskolin was then added (10 μM) on the apical side to activate CFTR, followed by the addition of Ivacaftor (10 μM) and/or 10 μM of AmB. The transepithelial electrical resistance was measured using the Acquire and Analyze 2.3 software.
